# Feature-specific awake reactivation in human V1 after visual training

**DOI:** 10.1101/231381

**Authors:** Ji Won Bang, Yuka Sasaki, Takeo Watanabe, Dobromir Rahnev

## Abstract

Converging human studies have demonstrated that brain activity patterns observed during task performance reemerge in the following restful awake state. Such “awake reactivation” has been demonstrated across higher-order cortex for complex images or associations. However, it remains unclear what specific training components are reactivated in these studies. Here we sought to provide evidence for the reactivation of a particular visual feature – Gabor orientation. Following extensive training on a visual task, we found robust reactivation in human V1 that lasted at least eight minutes. This effect was not present in higher retinotopic areas such as V2, V3, V3A, or V4v, demonstrating that the effects in V1 are not due to top-down processes such as conscious rehearsal. Furthermore, the amount of awake reactivation predicted the amount of performance improvement on the visual task. These results demonstrate that functionally-relevant awake reactivation of specific visual features occurs in early sensory cortex.

## Introduction

Understanding how the human brain learns is one of the central goals in neuroscience. A growing body of literature demonstrates that new learning becomes consolidated during offline states (Diekelmann & Born, 2010; McGaugh, 2000; Sasaki, Nanez, & Watanabe, 2010). More specifically, recent functional magnetic resonance imaging (fMRI) studies in humans point to memory reactivation – the reemergence of the brain activity patterns observed during task performance – as a potential contributor to offline memory consolidation (Deuker et al., 2013; Schlichting & Preston, 2014; Staresina, Alink, Kriegeskorte, & Henson, 2013; Tambini & Davachi, 2013). This reactivation occurs during sleep (Deuker et al., 2013) but also during restful wakefulness (Chelaru et al., 2016; de Voogd, Fernandez, & Hermans, 2016; Deuker et al., 2013; Guidotti, Del Gratta, Baldassarre, Romani, & Corbetta, 2015; Schlichting & Preston, 2014; Staresina et al., 2013; Tambini & Davachi, 2013) in which case it is referred to as *awake reactivation.*

Pioneering human studies have demonstrated the existence of awake reactivation for episodic memories in the medial temporal lobe including in the hippocampus (Tambini & Davachi, 2013), the entorhinal and the retrosplenial cortex (Staresina et al., 2013). Specifically, brain patterns during encoding of associations between different categories persisted into the following rest period (Staresina et al., 2013; Tambini & Davachi, 2013). The amount of reactivation in the hippocampus also predicted later behavioral performance (Tambini & Davachi, 2013).

Building on these early findings, several lines of human studies found that awake reactivation exists beyond the medial temporal lobe (Chelaru et al., 2016; de Voogd et al., 2016; Deuker et al., 2013; Guidotti et al., 2015; Schlichting & Preston, 2014). The brain patterns during encoding of associations between objects and spatial locations were shown to reemerge in both inferior temporal cortex and occipital cortex (Deuker et al., 2013). In addition, category-specific awake reactivation was demonstrated in the inferior temporal cortex (de Voogd et al., 2016) and the fusiform face area (Schlichting & Preston, 2014). More recent studies used diverse paradigms beyond hippocampus-dependent episodic memory. Repeated exposure to movies led to reactivation in a large cortical network, consisting mostly of the frontal and temporal lobes (Chelaru et al., 2016). Similarly, training on a letter discrimination task induced reactivation across medial and lateral parietal regions and higher-order visual cortex (Guidotti et al., 2015). This converging evidence suggests that awake reactivation occurs across higher-order areas beyond the medial temporal lobe.

However, previous research could not determine what specific training components were reactivated. Indeed, some previous studies used associative learning between different categories (Deuker et al., 2013; Schlichting & Preston, 2014; Staresina et al., 2013; Tambini & Davachi, 2013). The use of associative learning makes it difficult to disentangle brain processes related to the coding of each stimulus from the brain processes related to the binding of different stimuli (Deuker et al., 2013). The studies that did not use associative learning employed relatively complex visual stimuli such as animals, fruits, vegetables, letters and cartoon movies (Chelaru et al., 2016; de Voogd et al., 2016; Deuker et al., 2013; Guidotti et al., 2015; Schlichting & Preston, 2014; Staresina et al., 2013; Tambini & Davachi, 2013). These complex visual stimuli consist of a large number of individual features that could potentially be reactivated. Further, these studies observed awake reactivation in higher-order brain regions that have mixed selectivity to features (Huth, de Heer, Griffiths, Theunissen, & Gallant, 2016), making it again difficult to infer what specific features are reactivated.

Isolating a specific feature that is reactivated is critical for at least two reasons. First, it will demonstrate the limits of the phenomenon. Does reactivation in the human brain occur only at the level of complete objects? Is it a phenomenon that can be observed only in higher-level cortex? Or is awake reactivation a phenomenon that can be observed across the whole brain and at different levels? Second, providing evidence for awake reactivation of specific features will link the studies in humans even more tightly to the phenomenon of neuronal replay observed in animals where replay is typically demonstrated for a specific feature (e.g., location in space (Carr, Jadhav, & Frank, 2011; Davidson, Kloosterman, & Wilson, 2009; Diba & Buzsaki, 2007; Foster & Wilson, 2006) or placement on the screen (Han, Caporale, & Dan, 2008)).

Further, previous research often did not prevent subjects from consciously rehearsing the trained stimuli (de Voogd et al., 2016; Deuker et al., 2013; Guidotti et al., 2015; Schlichting & Preston, 2014; Tambini & Davachi, 2013). In fact, some of these studies may have even encouraged such conscious rehearsal. Therefore, it is unclear whether the observed awake reactivation in prior studies reflects spontaneous brain activity or purposeful, top-down effects. Since awake reactivation is usually conceptualized as an automatic process outside of conscious awareness, it is important that the phenomenon is demonstrated even when conscious rehearsal is explicitly prevented.

In the current study, we set to address above-mentioned limitations and provide evidence for reactivation of a specific visual feature that is not driven by conscious rehearsal. To do so, we focused on the primary visual cortex (V1), which is known to code stimulus orientation (Yacoub, Harel, & Ugurbil, 2008) and show changes after visual training (Rosenthal, Andrews, Antoniades, Kennard, & Soto, 2016; Sasaki et al., 2010). We used oriented Gabor patches rather than complex stimuli for training and examined whether the trained orientation is reactivated in V1 after extensive visual training. To prevent any conscious rehearsal, we required participants to perform a challenging fixation task while their pre- and post-training spontaneous brain activity was measured. Our results show that the activity patterns of V1 are more likely to be classified as the trained, compared to an untrained orientation shortly after visual training. However, higher retinotopic areas such as V2, V3, V3A, or ventral V4 (V4v), did not show this effect, thus providing direct evidence that the effects in V1 were not driven by top-down processes such as conscious rehearsal. Furthermore, greater amount of awake reactivation in V1 was found to be associated with greater learning amount on the trained stimulus. These findings demonstrate that feature-specific, spontaneous awake reactivation takes place after visual training in V1 and that this reactivation is functionally important for visual learning.

## Results

We examined whether a process of feature-specific, spontaneous awake reactivation occurs after visual training in human V1. We trained human subjects (n = 12) to detect a specific Gabor orientation (45° or 135°, counter-balanced between subjects; **Figure 1A**). The training induced significantly better performance on the post- compared to the pre-training test (F(1,11)=34.351, P<0.001; **Figure S1**), thus demonstrating that significant behavioral learning took place.

**Figure 1.**
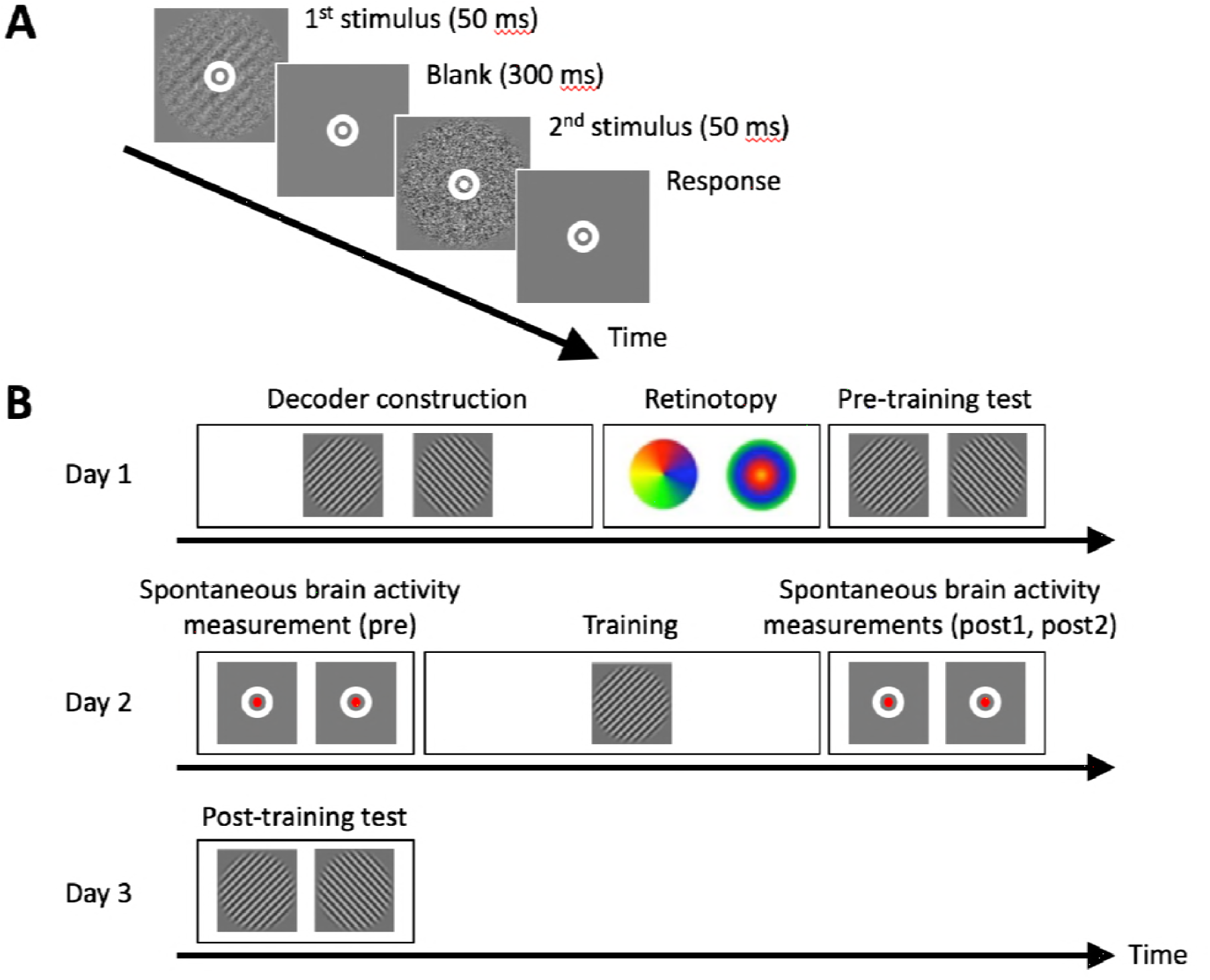
Task and experimental procedure. *(A) Subjects performed a 2-interval-forced-choice (2IFC) orientation detection task, in which they indicated whether a Gabor patch appeared in the first or second interval. (B) The experiment consisted of three days. Visual training (∼40 minutes on average) was conducted on Day 2. We collected two spontaneous measurement scans (5 minutes/scan) before (combined into a single pre baseline) and after training (post1 and post2). During these scans subjects detected a color change in a small fixation dot (see Methods). Decoder construction (∼50 minutes) and retinotopy (∼20 minutes) scans were collected on Day 1. Pre- and post-training behavioral tests were conducted on Days 1 and 3 to confirm that training improved task performance.*

Before the training, we constructed a decoder that could distinguish between the multivoxel pattern of blood-oxygen-level dependent (BOLD) activity elicited by each Gabor orientation (**Figure 1B**). We then applied this decoder on the spontaneous activity scans conducted before (pre, two 5-min scans that we combined for analysis) and after (post1 & post2: consecutive two 5-min scans analyzed separately) the training.

If feature-specific awake reactivation indeed occurs in V1, it would manifest itself as post-training, but not pre-training, spontaneous activity appearing more similar to the trained, compared to the untrained, orientation. To test for such feature-specific awake reactivation in V1, we analyzed the decoder’s probability of classifying spontaneous activity in V1 as the trained orientation. Note that in this analysis, spontaneous activity was always classified as either the trained or untrained Gabor orientation at each data point (see Methods), and therefore we only report the probability of “trained orientation” classifications. In order to examine whether the decoder’s classification for the trained orientation significantly increased after the training, we applied a one-way repeated measures ANOVA to the decoder’s probability. The result showed a significant main effect of time (pre vs. post1 vs. post2; F(2,22)=6.321, P=0.007). Consistent with the existence of feature-specific awake reactivation, the activation pattern in V1 was more likely to be classified as the trained orientation immediately after (post1) compared to before (pre) the visual training (t(11)=3.607, P=0.004, uncorrected paired sample t-test; **Figure 2A,B**). A significant quadratic trend was also observed in the time-course of the decoder’s probability (F(1,11)=12.041, P=0.005), indicating a peak right after training. Furthermore, the probability of classification as the trained orientation was significantly greater than chance in both post-training periods (post1: t(11)=4.467, P=0.001; post2: t(11)=2.929, P=0.014, one-sample t-tests) but not before training (pre, t(11)=0.243, P=0.812, one-sample t-test).

**Figure 2.**
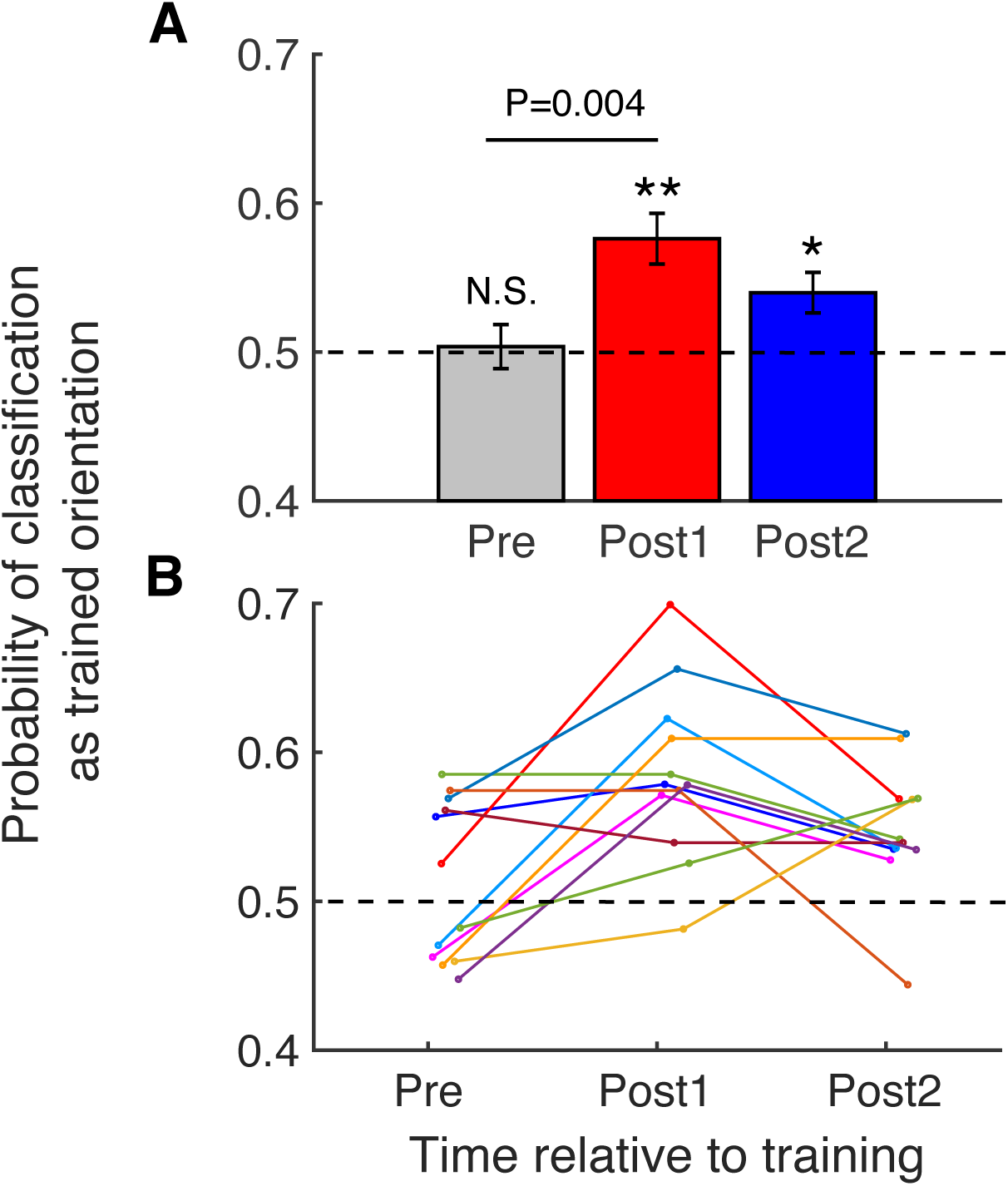
Probability of classifying spontaneous brain activity as trained orientation in V1. *Consistent with the existence of awake reactivation, spontaneous activity in V1 immediately after training was more likely to be classified as the trained orientation. The effect is visible both in the group (A) and individual data (B). Error bars represent s.e.m. * P<0.05, ** P<0.01, N.S. not significant.*

The above analysis focused on the decoder’s probability of classifying activity as the trained orientation based on the decoder’s binary classification (either trained or untrained orientation) at each data point. However, for each data point, the decoder could also produce a continuous measure: the likelihood that the data point came from the trained or untrained orientation. Analyzing these continuous likelihood values (rather than the binary classification performance), we still found the same pattern of results (main effect of time in one-way repeated measures ANOVA, F(2,22)=3.780, P=0.039; **Figure S2**; see Methods). The likelihood of classification as the trained orientation was also significantly greater than chance in both post-training periods (post1: t(11)=3.584, P=0.004; post2: t(11)=2.400, P=0.035, one-sample t-tests) but not before training (pre, t(11)=0.427, P=0.678, one-sample t-test). Thus, both methods of analysis show that posttraining spontaneous activity in V1 appears more similar to the trained orientation.

One potential concern is that these results could be due to subjects consciously imagining the trained stimulus during the spontaneous brain measurement (post1 and post2). We made such behavior unlikely by requiring subjects to perform a challenging task in which they monitored for a slight color change at fixation during the spontaneous brain measurement (**Figure 1B**; see Methods). We further delayed any post-training test until the next day, thus making it unnecessary for subjects to mentally practice the trained task immediately after the training was over.

Despite these precautions, it is still possible that some amount of conscious rehearsal of the trained stimulus took place. We reasoned that if any conscious rehearsal occurred, reactivation-like activity should be observed in higher visual areas since top-down processes such as visual imagery (Albers, Kok, Toni, Dijkerman, & de Lange, 2013), attention (R. B. Tootell et al., 1998), and working memory (Harrison & Tong, 2009) have at least as strong (and often much stronger) influence on these higher visual areas as on V1. We therefore analyzed the decoder’s probability of classifying spontaneous activity as the trained orientation based on activity patterns in V2, V3, V3A, and V4v. In order to test whether the decoder’s performance in higher visual areas changed after the training, we conducted a two-way repeated measures ANOVA with factors time (pre vs. post1 vs. post2) and region (V2, V3, V3A, and V4v). The ANOVA revealed no significant effects of time (F(2,22)=0.240, P=0.789), region (F(3,33)=2.113, P=0.117), or interaction between the two (F(6,66)=2.022, P=0.075). Despite the lack of significant overall effects (**Figure 3**; individual results are shown in **Figure S3**), we wanted to ensure that none of these higher retinotopic regions showed the pattern of results in V1 where decodability of the trained stimulus increased immediately after training (post1) compared to before training (pre). In fact, we found no such effects in V2, V3, V3A, or V4v (all P values > 0.6 for pre vs. post1 in each area before correction: V2, t(11)=0.333, P=0.746; V3, t(11)=-0.492, P=0.632; V3A, t(11)=-0.380, P=0.711; V4v, t(11)=-0.514, P=0.618, paired sample t-tests). Therefore, the results in V1 are unlikely to be due to subjects consciously rehearsing the trained stimulus.

**Figure 3.**
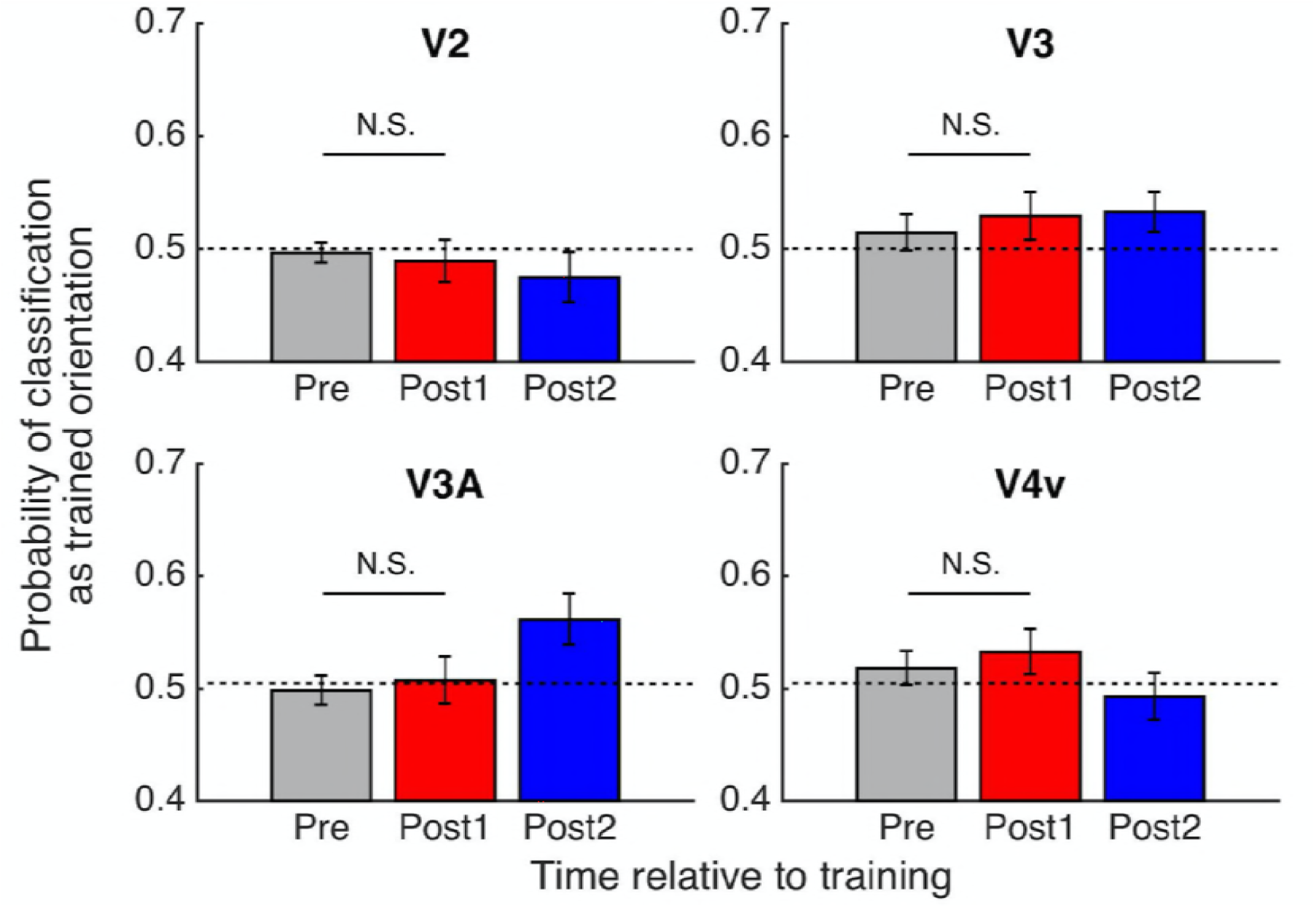
Probability of classifying spontaneous brain activity as trained orientation in V2, V3, V3A, and V4v. *Consistent with a lack of awake reactivation in higher retinotopic areas, spontaneous activity in V2, V3, V3A, and V4v was classified as the trained orientation at chance level (0.5) immediately after training. Error bars represent s.e.m. N.S. not significant.*

We analyzed the data from V1 separately from the other four retinotopic areas based on our a priori hypothesis. However, if all five retinotopic regions are entered into a twoway repeated measures ANOVA, then consistent with the existence of awake reactivation exclusively in V1, a significant interaction between region and time emerges (F(8,88)=2.303, P=0.027). In addition, we further examined whether the decoder’s probability of classifying spontaneous activity as the trained orientation in V1 is significantly greater than that in V2, V3, V3A, and V4v immediately after training (post1). We found a greater decodability of the trained stimulus in V1 compared to V2, V3A, and V4v immediately after training (post1) (V1 vs. V2, t(11)=4.196, P=0.001; V1 vs. V3, t(11)=1.478, P=0.168; V1 vs. V3A, t(11)=2.655, P=0.022; V1 vs. V4v, t(11)=2.708, P=0.020, paired sample t-tests, before correction). Again, these results support the notion that the pattern of increased decodability of the trained stimulus immediately after training is found exclusively in V1.

Finally, we examined whether the observed awake reactivation in V1 had functional consequences for learning. To address this question, we examined whether subjects who showed greater reactivation exhibited stronger behavioral improvement. We constructed subject-specific learning index that measures the amount of learning that was specific to the trained orientation. Similarly, we constructed a subject-specific reactivation index that measures the amount of reactivation by comparing the probability of decoding the trained stimulus in the pre and post-training spontaneous measurement scans (see Methods for details about both indices). We then compared the learning index across subjects with a high vs. low amount of reactivation (using a median split). We found that the learning index for those who showed higher awake reactivation was significantly greater than that for those who showed lower awake reactivation (t(10)=2.616, P=0.026, independent samples t-test; **Figure 4**). This result indicates that greater amount of awake reactivation is associated with greater visual learning on the trained stimulus and suggests that awake reactivation has a functional role in the learning.

**Figure 4.**
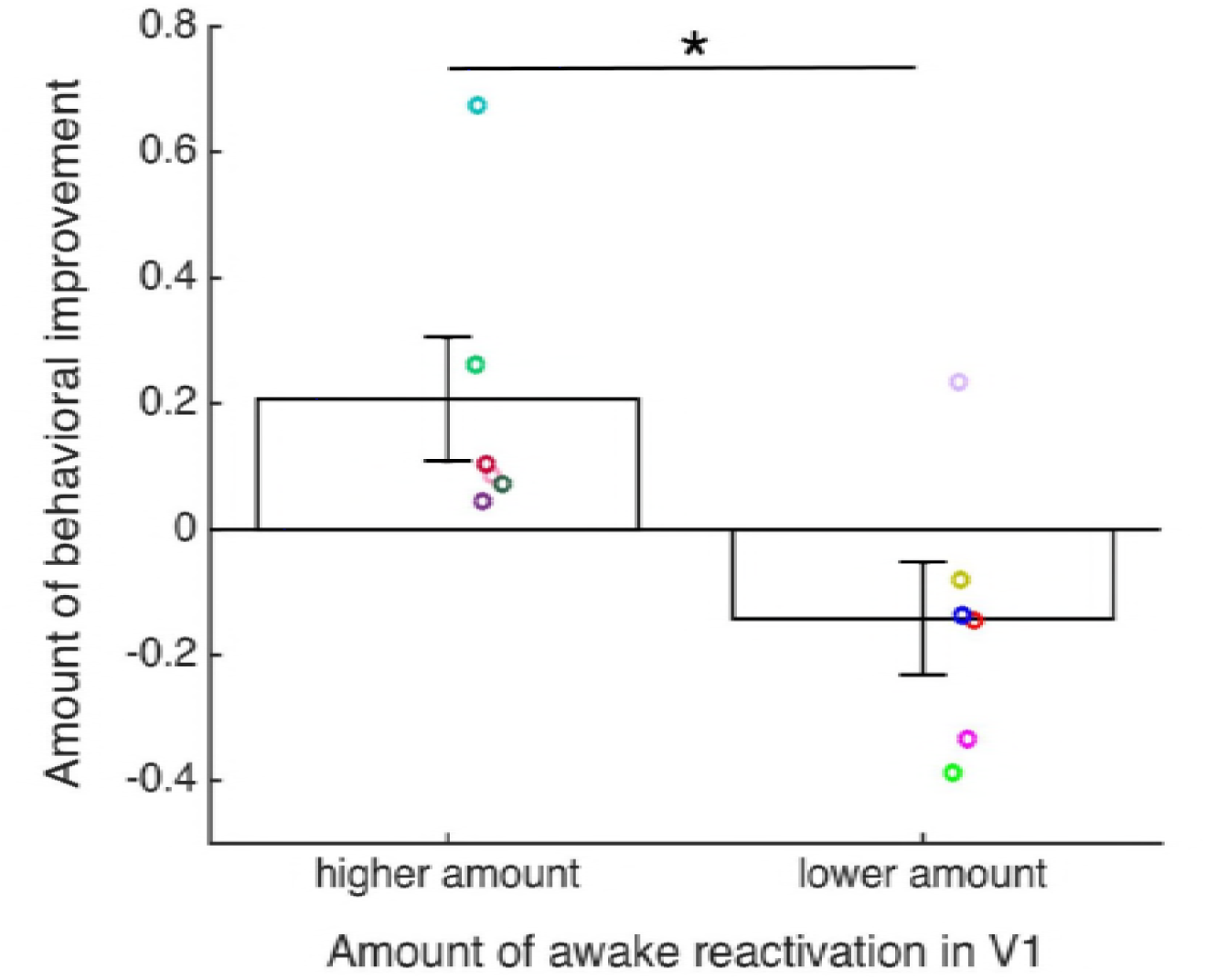
Strength of awake reactivation predicts performance improvement. *The amount of behavioral improvement specific to the trained orientation was significantly greater for subjects who showed higher levels of awake reactivation. Dots represent individual data. Error bars represent s.e.m. * P<0.05.*

## Discussion

We investigated whether a process of spontaneous, feature-specific awake reactivation takes place in human V1 following training with a novel visual stimulus. We found that shortly after training, the spontaneous brain activity in V1 was more likely to be classified as the trained orientation. This awake reactivation was specific to V1; higher retinotopic areas failed to show similar effects. Furthermore, the strength of awake reactivation predicted behavioral performance improvement on the trained stimulus. These results suggest that spontaneous, feature-specific awake reactivation occurs in V1 after the visual training and that this awake reactivation may serve to bolster the subsequent visual learning after the initial training is completed.

Our findings go beyond the previous literature on awake reactivation in three important ways. First, we demonstrate feature-specific reactivation. Indeed, by comparing Gabor stimuli that only vary in their orientation, we revealed that a basic visual feature such as orientation is specifically reactivated in the brain. Conversely, previous human studies could not determine what specific aspect of the trained stimulus was reactivated. Some of the prior studies used associative learning between two different categories (Deuker et al., 2013; Schlichting & Preston, 2014; Staresina et al., 2013; Tambini & Davachi, 2013). However, the reemergence of the brain processes during encoding of two different categories is insufficient to determine whether the reactivated information relates to the coding of each stimulus or the binding process (Deuker et al., 2013). Other studies did not use associative learning but employed complex stimuli such as animals, fruits, vegetables, letters and cartoon movies (Chelaru et al., 2016; de Voogd et al., 2016; Deuker et al., 2013; Guidotti et al., 2015; Schlichting & Preston, 2014; Staresina et al., 2013; Tambini & Davachi, 2013) that contain a large number of individual features that could potentially be reactivated. By demonstrating feature-specific reactivation, our study shows that awake reactivation in the human brain is not limited to the level of whole objects but extends down to the most basic visual features.

Second, we show that reactivation in humans is not exclusive to higher-order brain areas but exists even in the primary visual cortex V1. Prior studies demonstrated the existence of reactivation mostly in the association cortex but not in the primary sensory areas (Chelaru et al., 2016; de Voogd et al., 2016; Deuker et al., 2013; Guidotti et al., 2015; Schlichting & Preston, 2014; Staresina et al., 2013; Tambini & Davachi, 2013). Therefore, our study suggests that human reactivation is likely to occur across most (if not all) brain regions.

Third, our study demonstrates that awake reactivation can occur without conscious rehearsal. Most of the prior human studies on awake reactivation did not have a robust way of preventing subjects from engaging in conscious rehearsal (de Voogd et al., 2016; Deuker et al., 2013; Guidotti et al., 2015; Schlichting & Preston, 2014; Tambini & Davachi, 2013) and in some cases, may have actually encouraged such rehearsal. This issue is critical since awake reactivation is usually conceptualized as an automatic process outside of conscious awareness but evidence for this view has been lacking.

Several pieces of evidence suggest that awake reactivation in our study was not due to conscious rehearsal of the trained stimulus. First, we required subjects to perform a challenging task at fixation (see Methods) during the pre- and post-training periods. This fixation task introduced attentional demands that were not present in the no-task resting-state scans used in the previous studies. The presence of a demanding task made it unlikely that subjects would engage in a purposeful recall of the trained stimulus. At the same time, the fixation task was designed so that the parts of the visual cortex corresponding to the trained stimulus were not prevented from engaging in reactivation processes. This was achieved by making the target in the fixation task small and spatially non-overlapping with the trained stimulus. Second, we delayed any further tests on the trained task until the next day. This manipulation removed any immediate motivation to purposefully rehearse the trained stimulus during the fixation task. Third, we checked whether reactivation-like activity could be detected in higher retinotopic areas such as V2, V3, V3A, and V4v. It is known that top-down processes such as visual imagery (Albers et al., 2013), attention (R. B. Tootell et al., 1998), and working memory (Harrison & Tong, 2009) have a strong influence on these higher visual areas. The lack of reactivation in those higher regions thus shows that the results in V1 are indeed not due to conscious top-down rehearsal of the trained stimulus. Finally, after completing the experiment, we asked subjects if they mentally imagined the trained stimulus while performing the fixation task and they all denied doing so. Note, however, that a verbal statement alone should not be considered as sufficient evidence for a lack of conscious rehearsal.

Despite the fact that fMRI has limited spatio-temporal resolution compared to neuronal recordings, our results, together with prior human studies on awake reactivation, appear to reveal processes akin to neuronal replay in animals (Carr et al., 2011). Neuronal replay, that is the reemergence of neuronal patterns that represent previous learning, has been studied extensively in hippocampal cells (Carr et al., 2011; Davidson et al., 2009; Diba & Buzsaki, 2007; Foster & Wilson, 2006; Yao, Shi, Han, Gao, & Dan, 2007) but also in areas of the neocortex, particularly the visual (Han et al., 2008; Ji & Wilson, 2007; Yao et al., 2007) and the frontal cortex (Euston, Tatsuno, & McNaughton, 2007) in animals. More recent studies have shown that neuronal patterns representing previous learning can be preplayed by a cue even before a task begins (Eagleman & Dragoi, 2012; Ekman, Kok, & de Lange, 2017; Xu, Jiang, Poo, & Dan, 2012). These studies demonstrated the existence of cue-triggered preplay during wakefulness in both rodent V1 (Xu et al., 2012), monkey V4 (Eagleman & Dragoi, 2012) and human V1 (Ekman et al., 2017). Based on this converging evidence, we suspect that our results reflect spontaneous firing of the same pools of V1 neurons that code for the trained stimulus after training is completed. Future studies using finer spatio-temporal resolution are needed to directly examine this possibility.

In our study, the decodability of the trained orientation in V1 was highest in the first posttraining period (post1) but remained significant in the second post-training period (post2). Therefore, awake reactivation in our study continued at least until the onset of the second post-training period (post2), which occurred on average 7.75 minutes after the offset of visual training. In other words, reactivation persisted for at least ~8 minutes after subjects last experienced the trained stimulus. This time-course of awake reactivation in V1 agrees well with neuronal results in anesthetized rodent V1 (Han et al., 2008). In that study, 50 (125) stimulus repetitions led to ~3 (~14) minutes of replay activity. Compared to this study, our visual training was significantly longer but reactivation processes lasted for a shorter time. The shortened time length of reactivation is likely due to the fact that, unlike in anesthesia, the awake brain needs to remain responsive to its environment, which likely diminishes its capacity for reactivation. Future studies should include various training periods in order to delineate the time-course of the reactivation processes in humans.

In summary, we found evidence for spontaneous, feature-specific awake reactivation in area V1 after training on a visual task in humans. Critically, these effects were not observed in higher retinotopic areas. Further, the amount of awake reactivation in V1 predicted the behavioral learning amount on the trained stimulus. These results suggest that feature-specific awake reactivation might be a critical mechanism that supports offline memory consolidation.

## Materials and Methods

### Participants

Twelve healthy subjects (19 to 25 years old, 7 females) with normal or corrected-to-normal vision participated in this study. Exclusion criteria comprised a history of neurological, psychiatric disorders and contraindications to MRI. All subjects provided their demographic information and written informed consent, which was approved by the Institutional Review Board of Georgia Institute of Technology. All experiments were performed during daytime. All twelve subjects’ data were included in the analysis. Each experiment was performed once for all subjects.

### Procedures

The study consisted of three days (**Figure 1B**): decoder construction, retinotopy, and a pre-training test on Day 1, spontaneous brain activity measurements and training on Day 2, and a post-training test on Day 3. Days 1 and 2 were separated by multiple days, while Day 3 always immediately followed Day 2.

#### Orientation detection task

Subjects completed a 2-interval-forced-choice (2IFC) orientation detection task during training, as well as during the pre- and post-training tests. Subjects indicated which of two intervals contained a Gabor patch. The Gabor patch (contrast = 100%, spatial frequency = 1 cycle/degree, Gaussian filter sigma = 2.5 degree, random spatial phase) was presented within an annulus subtending 0.75 to 5 degrees and was masked by noise generated from a sinusoidal luminance distribution at a given signal-to-noise (S/N) ratio.

For instance, 10% S/N ratio meant that noise replaced 90% of the pixels in the Gabor patch. The interval without the Gabor patch consisted of pure noise (0% S/N ratio). The interval in which the Gabor patch was presented was determined randomly. During the entire orientation detection task, subjects were asked to fix their eyes on a white bull’s-eye on a gray disc (0.75° radius) at the center of the screen.

Each trial began with a 500-ms fixation period. After the fixation period, two stimuli were presented for 50 ms each, separated by a 300-ms blank period. Subjects indicated the interval in which the Gabor patch appeared by pressing a button on a keypad. No feedback was provided.

The training, as well as the pre- and post-training tests consisted of individual blocks. Within each block, we controlled the difficulty of the task by adjusting the S/N ratio using a 2-down 1-up staircase method. Each block started with stimuli at 25% S/N ratio and terminated after 10 reversals. Each block lasted approximately 1-2 minutes, contained 30-40 trials, and presented stimuli with just one orientation (45° or 135°).

#### Training

Training was conducted on Day 2 of our experiment. During the training, the subjects performed the 2IFC orientation detection task on one orientation only, randomly selected for each subject (to be either 45° or 135°). Subjects completed 16 blocks in total lasting approximately 40 minutes. The training was performed in the MR scanner.

#### Pre- and post-training tests

The purpose of the pre- and post-training tests was to obtain each subject’s threshold S/N ratio for each Gabor orientation. During these tests, subjects performed the same 2IFC orientation detection task. Each test consisted of two blocks, one for each Gabor orientation (45° or 135°). The order of presentation of the two orientations was randomized for each test and each subject. Subjects performed the pre- and post-training tests in a mock scanner located immediately adjacent to the scanner room.

#### Threshold S/N ratio

The threshold S/N ratio was determined separately for each block of testing. We computed the threshold S/N ratio as the geometric mean of the S/N ratio during the last 6 reversals in a block.

#### Relationship between awake reactivation and behavioral improvement

In order to check whether the amount of awake reactivation predicted the behavioral improvement, we first constructed an index of subjects’ learning. The learning index measures the relative learning amount for trained orientation compared to untrained orientation. The learning amount was defined as the change in the thresholds for pre- and post-training tests divided by the threshold at pre-training test:

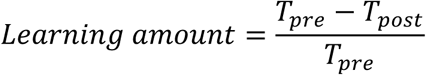

where *T_pre_* and *T_post_* refer to the threshold S/N ratios before and after training. We computed the learning index by subtracting learning amount for the untrained orientation from learning amount for the trained orientation.

To relate the learning index to the amount of awake reactivation, we constructed an equivalent index of subject-specific awake reactivation. The reactivation index measures the increase in decodability of the trained stimulus in V1 from the pre- to the post-training spontaneous brain measurement:

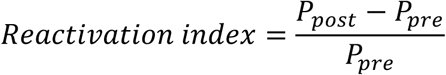

where *P_pre_* and *P_post_* refer to the probability of classifying brain activity as the trained orientation in the pre- and post-training spontaneous brain measurements, respectively.

#### Decoder construction

The purpose of the decoder construction stage was to obtain each subject’s blood-oxygen-level dependent (BOLD) signal patterns corresponding to each of the two Gabor orientations (45° and 135°). The data obtained from the decoder construction stage was later used to compute the parameters for the decoder that distinguishes between the two Gabor orientations.

During the decoder construction scan, the subjects performed 10 runs of frequency detection task (see below for details of the task) inside the MR scanner. Each run consisted of 18 trials, plus 6-sec fixation period in the beginning and the end of the run (1 run = 300 sec). Each trial had two parts, 12-sec stimulus presentation period and the following 4-sec response period (1 trial = 16 sec).

During 12-sec stimulus presentation period, 12 Gabor patches were presented at a rate of 1 Hz. Each Gabor patch was flashed for 500 ms allowing for 500 ms blank period between consecutive Gabor stimuli. All 12 Gabor patches had one specific orientation (45° or 135°) chosen randomly on each trial. In half of the 18 trials, one of the 12 Gabor patches had slightly higher spatial frequency compared to the other patches. For the other half of 18 trials, all 12 Gabor patches had the same spatial frequency. After the end of the 12-sec stimulus presentation, subjects indicated whether a Gabor patch with a different spatial frequency was presented. All Gabor patches were displayed with 50% S/N.

During each run, a white bull’s-eye on a gray disc (0.75° radius) was presented as a fixation point at the center of the screen. In the beginning of the 12-sec stimulus presentation period, the fixation point at the center of the screen changed its color from white to green to indicate that the stimulus presentation period had started. The color of the fixation point remained green throughout the 12-sec stimulus presentation period and changed to white again when the response period started.

The subjects were instructed to press a button on a key pad during 4-sec response period if they detected any change in the spatial frequency during preceding stimulus presentation period. If they did not detect any frequency change, they were instructed not to press a button. We controlled the difficulty of the task using an adaptive staircase method. The initial degree of the spatial frequency change in the first run was set to 0.24 Hz and decreased by 0.02 in the case of a hit and increased by 0.02 in the case of a false alarm or miss. In the case of a correct rejection, the degree of the frequency did not change. The following run always started from the last degree of the frequency in the previous run.

#### Retinotopy

We defined the retinotopically-organized areas V1, V2, V3, V3A, and ventral V4 (V4v) using standard retinotopic methods (Sereno et al., 1995; R. B. H. Tootell et al., 1997). Briefly, we presented a flickering checkerboard pattern at vertical/horizontal meridians and in the upper/lower visual fields. Based on the contrast map of BOLD signals, we delineated visual cortical areas of V1, V2, V3, V3A, and V4v. Furthermore, we presented an annulus stimulus in order to delineate the retinotopic regions in each visual area corresponding to the visual fields stimulated by the Gabor patch. The stimulus was a flickering checkerboard pattern within an annulus subtending 0.75 to 5 degree from the center of the screen, which is identical with the size of the Gabor patch. Only the voxels activated by the flickering pattern within an annulus stimulus were included in the main analyses.

#### Spontaneous brain activity measurement

We recorded each subject’s “spontaneous” BOLD activity before (pre, two 5-min scans that we combined for analysis) and after (post1 & post2: consecutive two 5-min scans analyzed separately) the visual training. In order to ensure that the subjects do not consciously imagine the trained orientation, we required them to perform a fixation task while they were scanned. During the fixation task, the subjects fixated their eyes on a white bull’s-eye on a gray disc (0.75° radius) at the center of the screen. They were instructed to press a button on a keypad as soon as they detected a color change of the center dot. A fixation dot changed its color from white ([R, G, B] = [255, 255, 255]) to faint pink ([R, G, B] = [255, 255 – x, 255 – x]) in an unpredictable way and returned to white 1.5 sec later. For the button press to be regarded as a hit, the subjects had to press a button within 1.5 sec (i.e., during the period when the color had changed). Initially, the color change x was set to 40 and then controlled by 2-down 1-up staircase rule.

To confirm that the subjects’ performance during the fixation task remained at a similar level over time (pre, post1, post2), we conducted a one-way repeated measures ANOVA on the accuracy rate with a factor of time (pre vs. post1 vs. post2). The result showed no significant main effect of time (F(2,22)=2.045, Huynh-Feldt correction, epsilon=0.724, P=0.169), suggesting that the subjects’ performance was constant throughout time.

### MRI Data Acquisition

Subjects were scanned inside a 3T MR scanner (Siemens 3T Trio) with a 12-channel head coil at Georgia Institute of Technology MRI Research Facility. For the anatomical reconstruction, high-resolution T1-weighted MR images were acquired using a multiecho magnetization-prepared rapid gradient echo (MPRAGE; 256 slices, voxel size = 1 × 1 × 1 mm^3^, TR = 2530 ms, FoV = 256 mm). Functional MR images were acquired using gradient echo EPI sequences (voxel size = 3 × 3 × 3.5 mm, TR = 2000 ms, TE = 30 ms, flip angle = 79°). Thirty-three contiguous slices were positioned parallel to the AC-PC plane to cover the whole brain.

### fMRI Data Analysis

We analyzed the brain data using Freesurfer software (http://surfer.nmr.mgh.harvard.edu/). Since we obtained each subject’s brain data on separate days (days 1 and 2), we processed the structural images from two different days with the longitudinal stream (Reuter, Schmansky, Rosas, & Fischl, 2012), that is known to create an unbiased within-subject structural template using robust, inverse consistent registration. All functional data were motion-corrected. No spatial or temporal smoothing was applied to the functional data. We registered the functional data from two different days’ scans to the individual structural template that was created via the longitudinal stream using rigid-body transformations. A gray matter mask was used for extracting BOLD signals from voxels located within the gray matter.

To localize the retinotopic areas V1, V2, V3, V3A, and V4v, as well as their sub-regions corresponding to the visual fields stimulated by the Gabor patch during the 2IFC orientation detection task, we performed a conventional amplitude analysis (see Retinotopy in Methods for more details about stimuli). Briefly, using the contrast map between vertical/horizontal meridians and upper/lower visual fields, we delineated visual cortical areas of V1, V2, V3, V3A, and V4v. Then using a separate scan during the Retinotopy session, we localized the sub-regions in each visual area corresponding to the part of the visual field occupied by the Gabor patch during the 2IFC orientation detection task. To do so, we contrasted two conditions: an annulus ON condition, in which a flickering checkerboard pattern filled an annulus subtending 0.75 to 5 degrees (same size as in the 2IFC task) and an annulus OFF condition in which a flickering checkerboard pattern filled the whole screen except an annulus subtending 0.75 to 5 degrees.

After we identified the sub-regions of V1, V2, V3, V3A, and V4v, we extracted the time-courses of BOLD signal intensities from each voxel within the sub-regions using MATLAB (MathWorks, Natick, MA). We shifted the BOLD signals by 6 sec to account for the hemodynamic delay and removed a linear trend. Then, within each run, we normalized (“z-scored”) each voxel’s BOLD time-courses to minimize the baseline changes across runs. Specifically, to create the data sample for decoding, we averaged the BOLD signals across 6 volumes (12 sec) that correspond to the duration of the stimulus presentation period in the decoder construction stage.

We used sparse logistic regression (SLR) (Yamashita, Sato, Yoshioka, Tong, &Kamitani, 2008) and the SLR toolbox (http://www.cns.atr.jp/~oyamashi/SLRWEB). SLR selects relevant voxels in the ROIs automatically while estimating their weight parameters for classification. We selected the voxels within the sub-regions of V1, V2, V3, V3A, and V4v as the input voxels. Then we trained the decoder to classify the BOLD patterns as either the Gabor stimulus with 45° or 135° orientation using 180 data samples from 10 runs of decoder construction stage (90 samples for each orientation). We tested the accuracy of the decoder using a leave-one-run-out cross-validation procedure for each of sub-regions of V1, V2, V3, V3A, and V4v. **Figure S4** shows the decoder accuracy for each ROI. The accuracy of the decoder was significantly above the baseline for all ROIs (all P values < 0.005).

Finally, we applied the same decoder to the spontaneous BOLD signals obtained before and after the visual training. The time-courses of the spontaneous BOLD signals were again averaged across 6 volumes. Then the decoder calculated how likely each spontaneous BOLD activity pattern averaged across 6 volumes is to the pattern elicited by the real Gabor orientation during decoder construction stage. Based on this likelihood result, the decoder classified each pattern averaged across 6 volumes to either trained or untrained orientation. **Figure 2, 3** and **Figure S3** show the probability of the brain activity being classified as the trained orientation. **Figure S2** shows the likelihood of the brain activity being classified as the trained orientation.

### Statistics

All statistical tests were two-tailed and the alpha level was set to 0.05. We used parametric statistical tests such as t-tests and ANOVA. For all repeated measures ANOVAs, we used Mauchly’s test of Sphericity to test the assumption of sphericity. We used Huynh-Feldt correction with the estimated epsilon when the sphericity assumption was violated. All such violations are reported when they occurred. All effects reported as significant remained such after application of Bonferroni correction. The sample size was determined based on similar fMRI experiments on visual learning (Guidotti et al., 2015; Shibata, Sasaki, Kawato, & Watanabe, 2016). The data collection and analysis were not conducted blindly by the experimenters.

### Apparatus

All visual stimuli were created via MATLAB and Psychtoolbox 3 on Mac OS X (Brainard, 1997). We presented visual stimuli on LCD display (1024 × 768 resolution, 60 Hz refresh rate) inside a mock scanner and via MRI-compatible LCD projector (1024 × 768 resolution, 60 Hz refresh rate) inside a 3T MR scanner.

### Data and Software Availability

The data and the computer codes are freely available online at: https://osf.io/9du8v/

## Acknowledgements

This work was funded by a startup grant to D.R. from the Georgia Institute of Technology.

## Author Contributions

J.W.B., Y.S., T.W. and D.R. conceived the study; J.W.B. and D.R. collected and analyzed data; J.W.B., Y.S., T.W. and D.R. wrote the manuscript.

---

**Figure S1.**
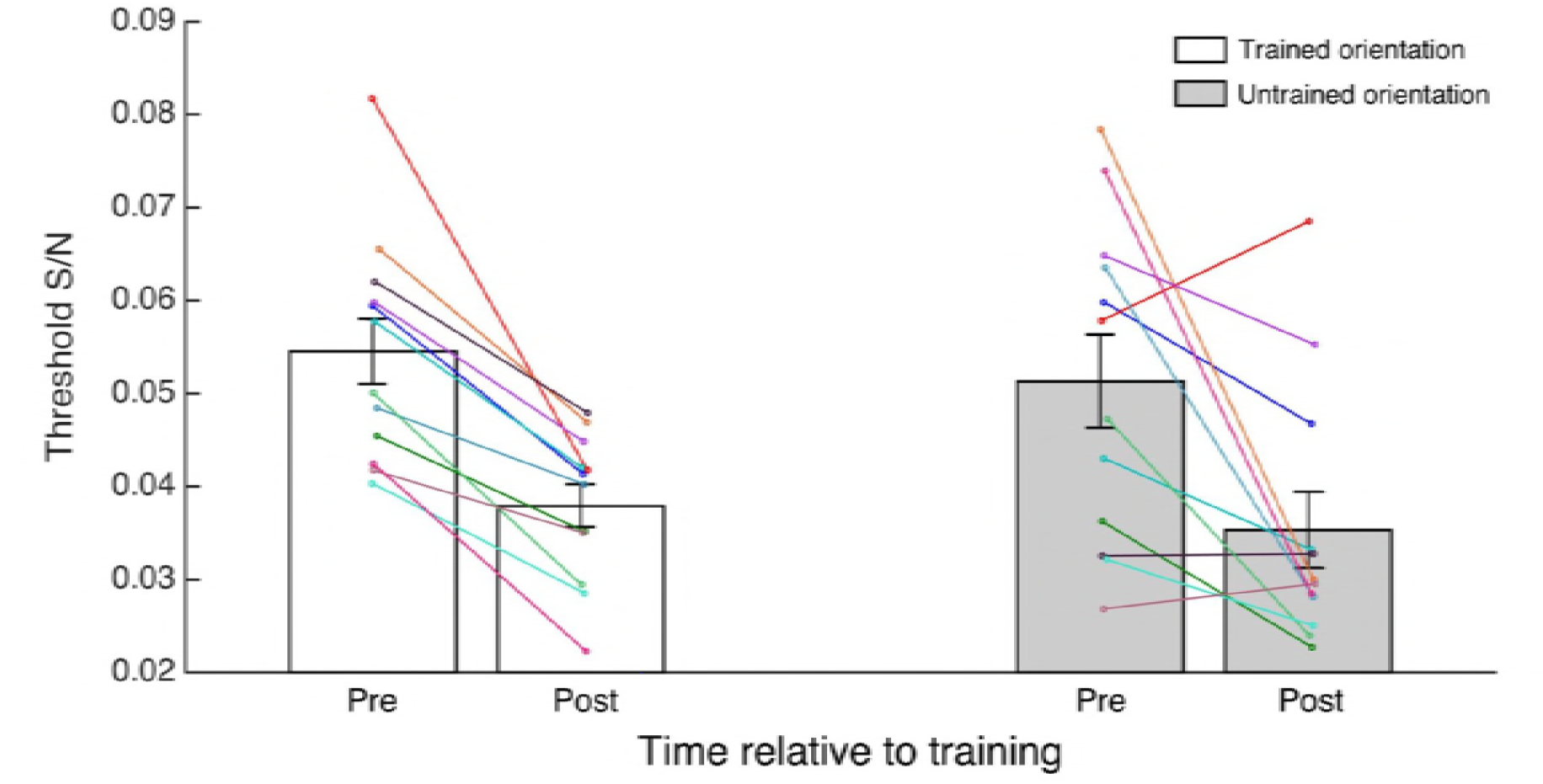
Threshold S/N (mean ± s.e.m.) during pre- and post-training stages for trained (white) and untrained (gray) orientations. To confirm that the subjects showed learning between pre- and post-training test stages, we performed a two-way repeated measures ANOVA with factors of orientation (trained vs. untrained orientation) and time (pre- vs. post-training) on the threshold S/N ratio. The results showed a significant main effect of time, indicating that the learning occurred for both orientations (time effect: F(1,11)=34.351, P<0.001). The dots represent individual data.

**Figure S2.**
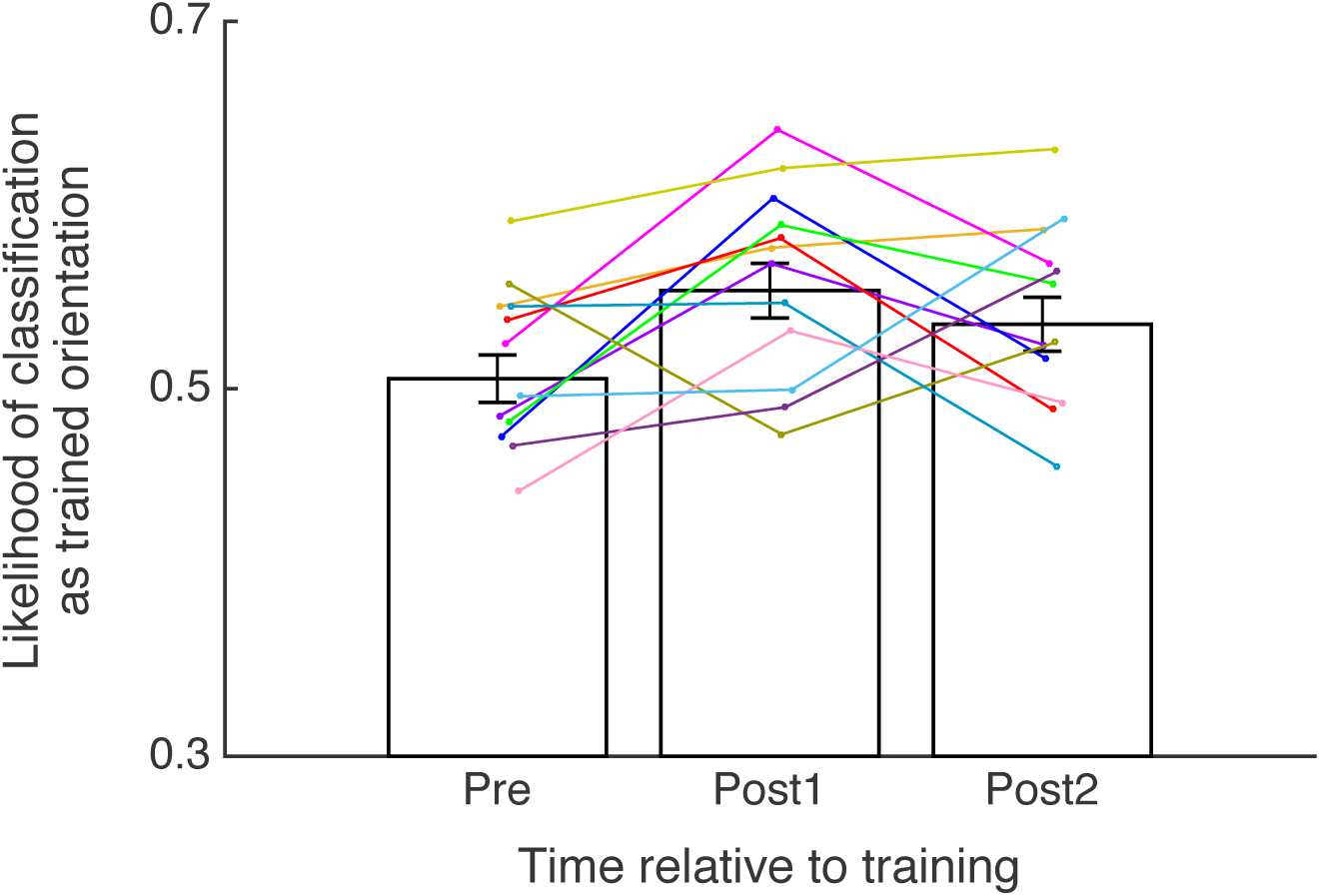
Likelihood of classification as trained orientation in V1. Individual data is plotted on the group data (mean ± s.e.m.). The likelihood of classification as trained orientation refers to a continuous measure (rather than a binary decision) of likelihood that a particular brain signal corresponds to the trained or untrained orientation. A oneway repeated measures ANOVA to the decoder’s likelihood of classification as the trained orientation showed a significant main effect of time (pre vs. post1 vs. post2; F(2,22)=3.780, P=0.039). The likelihood of classification as the trained orientation increased immediately after (post1) compared to before (pre) the visual training (t(11)=- 2.750, P=0.019, uncorrected paired sample t-test). In addition, the likelihood of classification as the trained orientation was significantly higher than chance immediately after training (post1, t(11)=3.584, P=0.004, one-sample t-test) and 5 minutes after training (post2, t(11)=2.400, P=0.035, one-sample t-test) but not before training (pre, t(11)=0.427, P=0.678, one-sample t-test). Consistent with the result from the probability of classification, this result indicates that post-training spontaneous activity in V1 appears more similar to the trained orientation.

**Figure S3.**
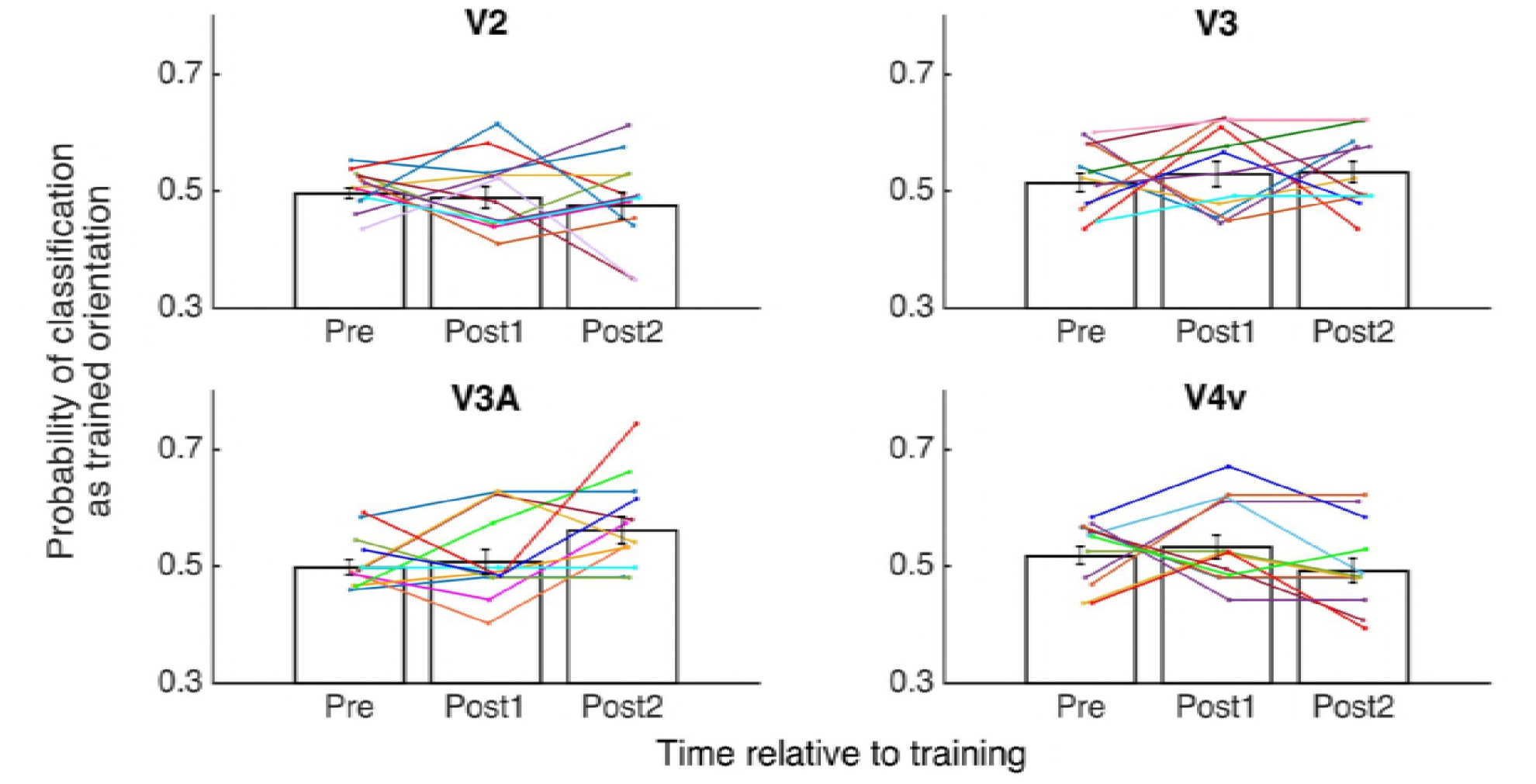
Probability of classifying spontaneous brain activity as trained orientation in V2, V3, V3A, and V4v. Individual data is plotted on the group data (mean ± s.e.m.) for each region of interest. The probability for the trained orientation did not increase immediately after training in V2, V3, V3A, and V4v.

**Figure S4.**
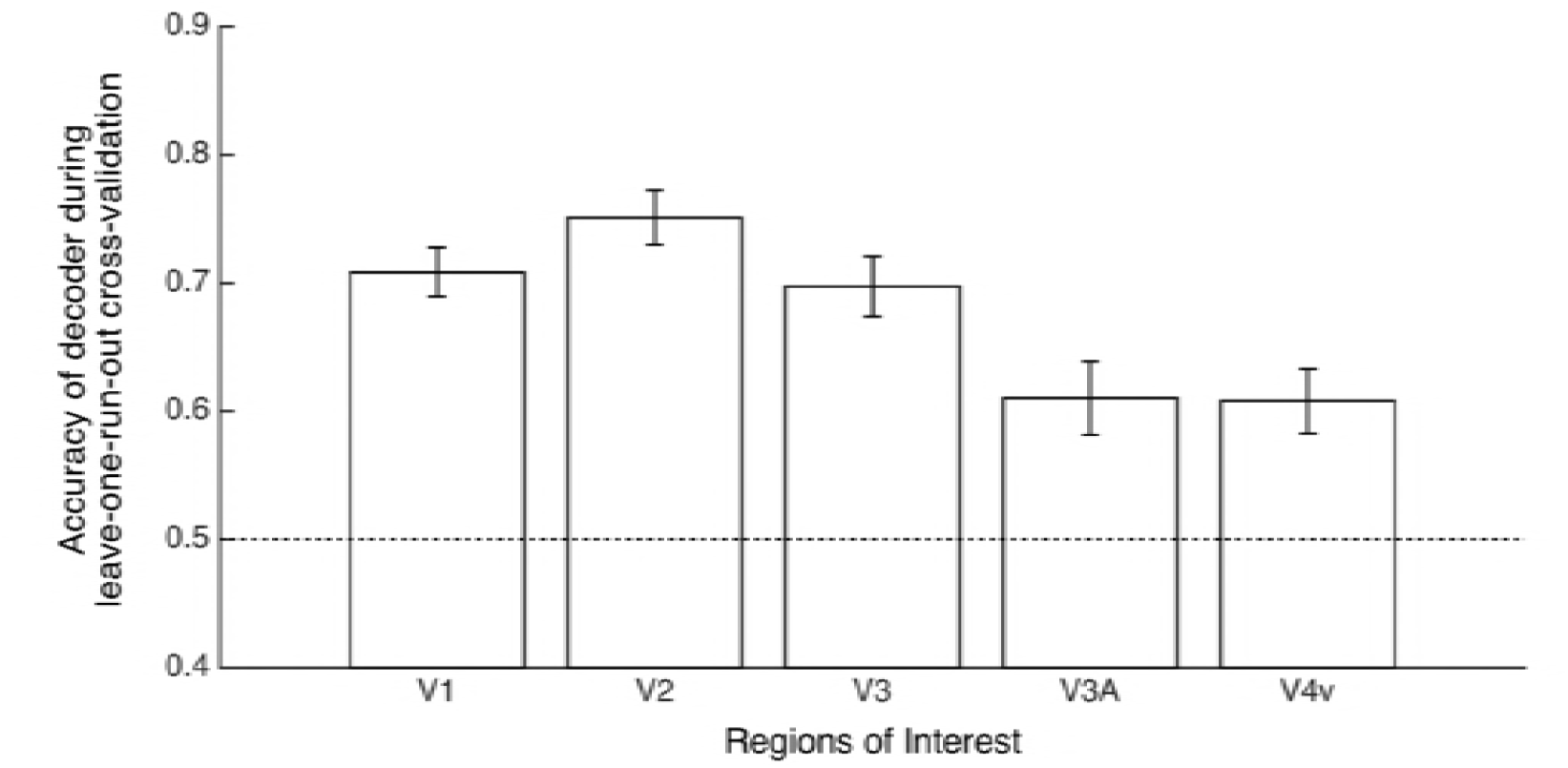
Accuracy of decoder during leave-one-run-out cross-validation in V1, V2, V3, V3A and V4v. The decoder was tested through ‘leave-one-run-out’ cross-validation using brain data obtained from the decoder construction scan. The accuracy of the decoder (mean ± s.e.m.) was above chance level (0.5) for all regions of interest (all P values < 0.005 before correction, one-sample t-tests; V1, t(11)=10.856, P=0.000; V2, t(11)=11.746, P=0.000; V3, t(11)=8.345, P=0.000; V3A, t(11)=3.851, P=0.003; V4v, t(11)=4.241, P=0.001).

